# Action selection based on multiple-stimulus aspects in the wind-elicited escape behavior of crickets

**DOI:** 10.1101/2021.04.23.441064

**Authors:** Nodoka Sato, Hisashi Shidara, Hiroto Ogawa

## Abstract

Animals detect approaching predators via sensory inputs through various modalities and immediately show an appropriate behavioral response to survive. Escape behavior is essential to avoid the predator’s attack and is more frequently observed than other defensive behaviors. In some species, multiple escape responses are exhibited with different movements. It has been reported that the approaching speed of a predator is important in choosing which escape action to take among the multiple responses. However, it is unknown whether other aspects of sensory stimuli, that indicate the predator’s approach, affect the selection of escape responses. We focused on two distinct escape responses (running and jumping) to a stimulus (short airflow) in crickets and examined the effects of multiple stimulus aspects (including the angle, velocity, and duration) on the choice between these escape responses. We found that the faster and longer the airflow, the more frequently the crickets jumped, meaning that they could choose their escape response depending on both velocity and duration of the stimulus. This result suggests that the neural basis for choosing escape responses includes the integration process of multiple stimulus parameters. It was also found that the moving speed and distance changed depending on the stimulus velocity and duration during running but not during jumping, suggesting higher adaptability of the running escape. In contrast, the movement direction was accurately controlled regardless of the stimulus parameters in both responses. The escape direction depended only on stimulus orientation, but not on velocity and duration.

**Summary statement:** When air currents triggering escape are faster and longer, crickets more frequently jump than run. Running speed and distance depend on stimulus velocity and duration, but direction control is independent.

## INTRODUCTION

Selecting a behavioral response appropriate to a situation is crucial for animals to survive events such as predator attacks. Vertebrates and invertebrates universally exhibit escape behavior to survive these attacks (Domenici et al.,2011a, b; LeDoux and Daw, 2018). Usually, animals quickly move away from the predator as a typical escape response, which increases their survival rate. The more emergent the situation is, the more predominantly the escape response is chosen compared to other defensive behaviors such as freezing (Baba and Shimozawa, 1997; Fadok et al., 2017; Evans et al., 2018; Turner et al., 2016). For instance, mice exhibit either a flight or freeze in response to the movement of the predator (De Franceschi et al., 2016); they choose the flight response when raptors approach them but exhibit freezing against raptors that just cruise past. This is because when the predator is unaware of the prey, freezing is more efficient to avoid being targeted. In contrast, once the predator is aware of the prey and begins to approach it, flight is an appropriate response to escape from the attack (Eilam, 2005). Since the animals of prey are subjected to attacks from various species of predators (Bulbert et al., 2015; Evans et al., 2019) and the approach of the predator varies trial by trial (Dangles et al, 2006; Walker et al., 2005), it is critical for the animals of prey to know how the predator approaches them, because it helps them make the appropriate behavioral choices against the emergency attack.

Animals detect the predator’s approach through various parameters of sensory stimuli, such as speed, duration, and orientation (Domenici, 2010). For instance, stimuli with a higher velocity indicate a larger predator and quicker attack, and those with longer duration suggest greater movement of the predator (Casas and Dangles, 2010). The orientation of the stimulus also indicates the direction from which the predator approaches. Based on these characteristics of the stimuli, the prey regulates its escape response. The stimulus velocity and orientation are related to the escape distance and direction, respectively (Bhattacharyya et al., 2017; Card, 2012; Domenici et al., 2011a, b; Dunn et al., 2016; Stewart et al., 2013). The size of the looming stimulus indicates the distance of the approaching predator and has been known to determine escape initiation in several species (Bhattacharyya et al., 2017; Dunn et al., 2016; Fotowat and Gabbiani, 2007; Preuss et al., 2006). The escape behavior of various animals employ multiple types of responses with distinct movements (Briggman et al., 2005; Liu and Hale, 2017; Takagi et al., 2017), resulting in different locomotor performance. Each escape response is considered to have a different advantage in performance, such as speed or controllability, which affect the success rate of avoidance from the predator’s attack (Card and Dickinson, 2008; Sato et al., 2019). Recent studies suggest that the approaching speed of the predator indicated by the stimulus velocity has an impact on the action selection between different escape responses, such as short and long modes of flies taking-off (Card and Dickinson, 2008; von Reyn et al., 2014, 2017). However, it is unknown what other parameters, including the direction or duration of the stimulus, are involved in the action selection for escaping.

We addressed this issue using the wind-elicited escape behavior of the field cricket *Gryllus bimaculatus* as the experimental material. Crickets exhibit either running or jumping in response to a short air current that is detected as a predator’s approach (Dupuy et al., 2011; Sato et al., 2017, 2019). The running and jumping movements are different from each other: the crickets either run on the ground or jump by strongly kicking the ground. Our previous study has revealed a “trade-off” between speed and behavioral flexibility: if crickets choose to run rather than jump, they move more slowly but can respond more flexibly to further attacks received during the response (Sato, et al., 2019). Interestingly, crickets can also control their movement direction accurately in response to the stimulus from any direction, not only during running but also while jumping (Sato et al., 2019), in contrast to escape by jumping in locusts (Santer et al., 2005; Simmons et al., 2010). Besides, the longer the stimulus, the longer the distance the crickets run (Oe and Ogawa, 2013). These facts suggest that crickets may be able to take multiple stimulus parameters into account while choosing either to run or jump.

In this study, we manipulated the three parameters of an air-current stimulus, (angle, velocity, and duration) and examined their effects on the response probabilities to run and jump those were elicited by the stimulus. Furthermore, to clarify the impacts of these parameters on the performance in the escape responses, we examined the relationships between the locomotor parameters, which included movement speed, travel distance, reaction time, directionality, and other stimulus parameters. Our results demonstrate that crickets choose the escape response and regulate their locomotion based on multiple stimulus parameters.

## MATERIALS AND METHODS

### Animals

We used the wild-type strain of crickets (*Gryllus bimaculatus*, Hokudai WT; Watanabe et al., 2018) that were bred in our laboratory. Thirty adult male crickets, less than 14 days after adult molting, were used throughout the experiments. They were reared under 12/12 h light/dark conditions at a constant temperature of 27°C. All experiments were conducted during the dark phase at 26-28°C.

### Behavioral experiment

The experimental apparatus as per our previous study (Sato et al., 2019) was used throughout the experiments. We monitored the wind-elicited escape responses using a high-speed digital camera (CH130EX, Shodensha, Osaka, Japan) installed above a circular arena (ø = 260 mm). After anesthetization by cooling with ice (0 °C) for 10 min, crickets were marked with two white spots on the dorsal surface of the head and thorax, the size of which was large enough to detect the movement of crickets from images. All experiments were performed after 30 min to make sure they completely recover from anesthetization. The cricket was placed in the center of the arena inside an inverted beaker (ø = 50 mm) covered with aluminum foil. After the beaker was carefully lifted, an air-current stimulus was immediately applied to the cricket that was standing still, and the response was recorded by the camera. Based on the video data (shutter speed, 1 ms; sampling rate, 120 frames s^−1^; total recording duration, 1660 ms), the two markers on the animal were automatically traced, and locomotor parameters mentioned in a later section were measured using motion analysis software (Move-tr/2D, Library, Tokyo, Japan). To monitor the entire trajectory during movement, we adopted a 285.7 × 285.7 mm frame size with 1024 × 1024 pixels resolution, which covered the entire arena.

### Stimulation and experimental procedure

The air-current stimulus used throughout the experiments was a short puff of nitrogen gas from a plastic nozzle (ø = 15 mm) connected to a pneumatic picopump (PV820, World Precision Instruments, Sarasota, FL, USA). One air-current nozzle was installed on the inside wall of the arena to be positioned on the same horizontal plane as the animal. Since the crickets were oriented randomly within a beaker, the stimulus angles measured as the angles of the crickets’ orientation against the stimulus source (left in Fig. 1A) were varied across the trials.

**Fig. 1.**
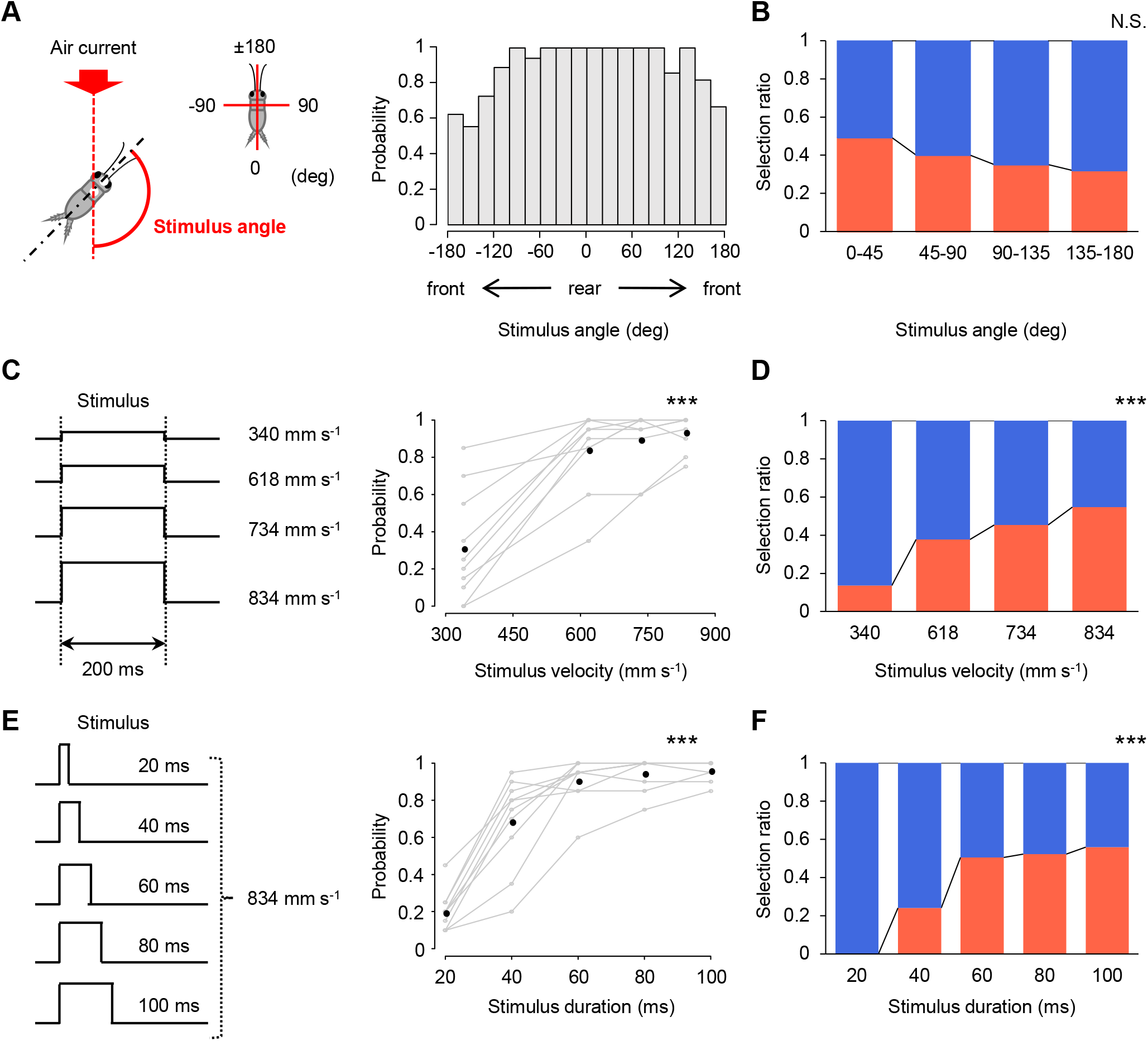
Effects of three stimulus parameters on action selection of running and jumping. **(A)** Distribution of the response probability against the stimulus angle in the angle test. Left diagram shows the definition of the stimulus angle. Right histograms represent the ratio of the number of responses to the number of trials for a range of every 20 degree of the stimulus angles. The air current with a velocity of 834 mm s^−1^ and a duration of 200 ms was used. *N*=10 animals. **(B)** The selection ratio of running (blue) to jumping (red) in the angle test. The data is shown in four divisions based on the absolute value of the stimulus angle. **(C)** Changes in the response probability over the stimuli with different velocities. Left diagram indicates four types of stimuli used in the velocity test, in which velocity was varied and duration was fixed to 200 ms. *N*=10 animals. **(D)** The selection ratio of running (blue) to jumping (red) in the velocity test. **(E)** Changes in the response probability over the stimuli with different durations. Left diagram indicates five types of stimuli used in the duration test, in which duration was varied and velocity was fixed to 834 mm s^−1^. *N*=10 animals. **(F)** The selection ratio of running (blue) to jumping (red) in the duration test. In (C) and (E), gray open circles connected with gray lines represent the probability for each individual, and black filled circles represent the mean of the individuals’ probability, for each velocity or duration of stimuli. \*\*\**P*<0.001, N.S. not significant, coefficients for each of the stimulus parameter in logistic regression analysis.

The data of three experiments with different stimulation were analyzed to test the effect of the three stimulus parameters on escape responses: “angle test,” “velocity test” and “duration test.” A part of the data obtained in our previous study (Sato et al., 2019) was also analyzed for the angle test, in which the effect of stimulus angles was examined in the escape responses to a constant stimulus duration of 200 ms and velocity of 834 mm s^−1^. The travel time of the air current to the center of the arena was 14.6 ± 0.1 ms, which was calculated based on the delay in stimulus-evoked ascending spikes of projection neurons that were extracellularly recorded (*N* = 2 animals, 10 trials in total). In the velocity test, we examined the effect of stimulus velocities on the escape response to the stimuli of 200 ms duration with four different velocities (340, 618, 734, and 834 mm s^−1^) (left in Fig. 1C). The travel times of these stimuli were 21.2 ± 1.0, 15.5 ± 0.1, 14.6 ± 0.1 and 14.6 ± 0.1 ms for 340, 618, 734 and 834 mm s^−1^ of air currents, respectively (*N*=2 animals, 10 trials in total for each stimulus velocity). In the duration test, we examined the effect of stimulus duration on the escape response to the stimuli of 834 mm s^−1^ for four different durations (20, 40, 60, 80, and 100 ms) (left in Fig. 1E). The velocity and duration of the air current were regulated by air pressure and using the interval between the opening and closing of the valve of the picopump.

In the angle test, 20 trials were repeatedly performed for each individual at inter-trial intervals of 60–90 s. In both the velocity and duration tests, stimuli with different velocities or durations were applied to the same individuals at increasing (*N*=5 animals) or decreasing order (*N*=5 animals). Twenty trials were performed in one session for each type of stimulus, so in total, 80 or 100 trials were performed for each individual in tests. The crickets were rested between the sessions for approximately 10 min within a plastic container (138 mm × 220 mm × 135 mm) and freely accessed food and water. Ten crickets were tested for each experiment.

### Data analysis

The wind-elicited responses were analyzed as in our previous study (Sato et al., 2017, 2019). Responses were defined based on the translational velocity of the cricket movement. If the velocity exceeded 10 mm s^−1^ within 250 ms after the stimulus onset and was greater in its maximum value by more than 50 mm s^−1^, the cricket was considered to respond to the stimulus. If the cricket did not begin to move within 250 ms of the response definition period, the trial was considered as “no response.” For the angle test, the data in all trials were pooled, and then the response probability was calculated from the number of responding trials per total trials for the range of every 20 degree of stimulus angles (right in Fig. 1A). In the velocity and duration tests, the response probability was calculated from the number of responding trials per 20 trials for each type of stimulus (right in Fig. 1C,E).

As in our previous study (Sato et al., 2019), the wind-elicited responses were categorized into “running” or “jumping” according to the movement of legs during the locomotion, which was confirmed visually for all responding trials by frame check of the video. If all six legs were off the ground simultaneously, the response was defined as “jumping,” if any one of the six legs touched the ground during movement, that response was defined as “running” (Fig. S1A). We rarely observed complex behaviors combined with running and jumping, such as jumping after running and *vice versa*. The selection ratio of running or jumping was calculated as the proportion of responses for all responding trials. In the angle test, all trials were pooled and divided into four groups based on the range of absolute stimulus angle, 0–45, 45–90, 90–135, and 135–180, and the selection ratio was calculated for each angular range (Fig. 1B). In the velocity and duration tests, the selection ratio was calculated for each type of stimulus, regardless of the stimulus angle (Fig. 1D,F).

We focused on the “initial response” in the responding trial and analyzed the cricket’s movement during the initial response as defined in our previous studies (Fukutomi et al., 2015; Oe and Ogawa, 2013; Sato et al., 2017, 2019). The initial response was the first continuous movement (bout) in which the translational velocity was greater in its maximum value than 50 mm s^−1^ and never less than 10 mm s^−1^ after the stimulus onset. Therefore, the response start was measured as the time when the translational velocity exceeded 10 mm s^−1^ after stimulus onset and the response finish was measured as the time when the velocity was less than 10 mm s^−1^. Movement distance, maximum translational velocity, and reaction time were measured as the metric locomotor parameters that characterized the response movements. The movement distance was measured as the entire path length of the moving trajectory. The reaction time was calculated by subtracting the mean travel time of the air current, as mentioned above, from the time from the opening of the picopump to the start of the response. Angular parameters including movement direction and turn angle were calculated based on the cricket’s body axis, as a vector connecting the thoracic and head markers (Fig. S1B). The movement direction was measured as the angle between the body axis at the starting point of the response and a line connecting the thoracic markers at the start and finish points. Thus, if the cricket moved in the direction opposite to the stimulus source, the movement direction would be equal to the stimulus angle. Since it has been confirmed in a previous study that crickets move in the opposite direction to the stimulus in both running and jumping (Sato et al., 2019), the accuracy in the directional control was assessed by the absolute value of the difference between the movement direction and stimulus angle. The turn angle was measured as the angle between the body axes at the start and finish points. If the cricket is oriented to the exact opposite side of the stimulus source at the finish points, the turn angle would be equal to the stimulus angle.

### Statistical analysis

R programming software (ver. 3.4.4, R Development Core Team) was used for all the statistical analyses. We assessed the effect of the stimulus parameters on the response probability and selection ratio, both of which were calculated for each individual, using multiple logistic regression analysis in each of the angle, velocity, and duration tests. In addition to the manipulated stimulus parameter, the trial order was considered as an explanatory variable, as follows:

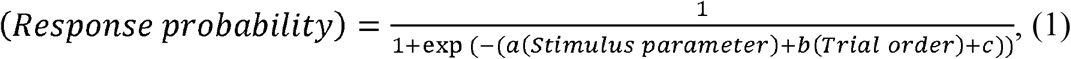

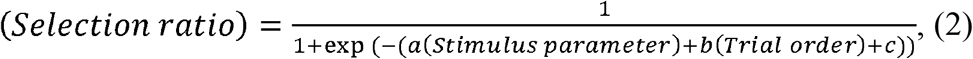

where *a* and *b* are the estimated coefficients, and *c* is the intercept. Then, the significance of the coefficients for stimulus parameters as explanatory variables was checked to assess their effect on the response probability or selection ratio (Table 1).

**Table 1.** Results of multiple logistic regression analysis to test the effect of stimulus parameters and trial order on the response probability and selected ratio.

| Experiment | Parameter | Explanatory variable | Result of logistic regression |  |  |
| --- | --- | --- | --- | --- | --- |
|  |  |  | Coefficient | z value | p value |
| “angle test” | Selection ratio | Stimulus angle | -0.269 | -1.841 | 0.066 |
|  | (Fig. 1B) | Trial order | -0.002 | -0.083 | 0.934 |
| “velocity test” | All probability | Stimulus velocity | 0.008 | 13.81 | <0.001 |
|  | (Fig. 1C) | Trial order | -0.015 | -0.882 | 0.378 |
|  | Selection ratio | Stimulus velocity | 0.004 | 5.896 | <0.001 |
|  | (Fig. 1D) | Trial order | 0.003 | 0.185 | 0.853 |
| “duration test” | All probability | Stimulus duration | 0.072 | 14.57 | <0.001 |
|  | (Fig. 1D) | Trial order | -0.014 | -0.865 | 0.387 |
|  | Selection ratio | Stimulus duration | 0.023 | 6.722 | <0.001 |
|  | (Fig. 1F) | Trial order | 0.006 | 0.473 | 0.636 |

To analyze the relationships between the metric locomotor parameters, such as movement distance, maximum velocity, or reaction time, and the stimulus angle, which was indicated as an absolute value in degrees from 0 (front) to 90 (lateral) and 180 (behind), we used linear regression analysis as follows:

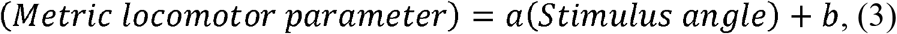

where *a* is the estimated coefficient and *b* is the intercept. The significance of the coefficients was checked to assess the effect of the stimulus angle on the metric parameters (Table 2).

**Table 2.** Results of linear regression analysis to test the relationships between the metric locomotor parameters and stimulus angle.

| Parameter | Response | Result of linear regression |  |  |
| --- | --- | --- | --- | --- |
|  |  | Estimated slope | t value | p value |
| Movement distance<br>(Fig. 2A) | Running | -0.104 | -3.346 | 0.001 |
|  | Jumping | -0.073 | -1.213 | 0.229 |
| Max. velocity<br>(Fig. 2B) | Running | -0.360 | -2.160 | 0.033 |
|  | Jumping | 0.652 | 2.105 | 0.039 |
| Reaction time<br>(Fig. 2C) | Running | 0.140 | 3.076 | 0.003 |
|  | Jumping | 0.021 | 0.545 | 0.587 |

Since the response probability and selection ratio were strongly affected by stimulus velocity and duration, the sample size of data for the locomotor parameters varied among the sessions using different types of stimuli. In particular, slow (340 mm s^−1^ of velocity) or short (20 ms of duration) air current rarely or never elicited jumping; therefore, we could not calculate the mean value of the locomotor parameters for each tested individual in the session using such stimuli. Considering the small and varied sample sizes, a linear mixed-effects model (LMM, the package ‘lme4’ ver. 1.1-23 in R) was adopted to assess the effects of stimulus velocity and duration on the locomotor parameters. We assumed the LMMs with stimulus parameters as explanatory variables, as follows:

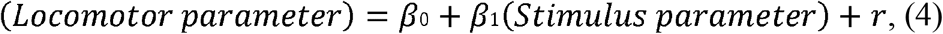

where *β*_0_ is the intercept, *β*_1_ is the estimated coefficient, and *r* is the random effect that the animal IDs were considered. The significance of the effect of stimulus parameters was tested by comparing the LMMs with and without the explanatory variable of stimulus velocity or duration by the likelihood ratio test (Table 3).

**Table 3.**
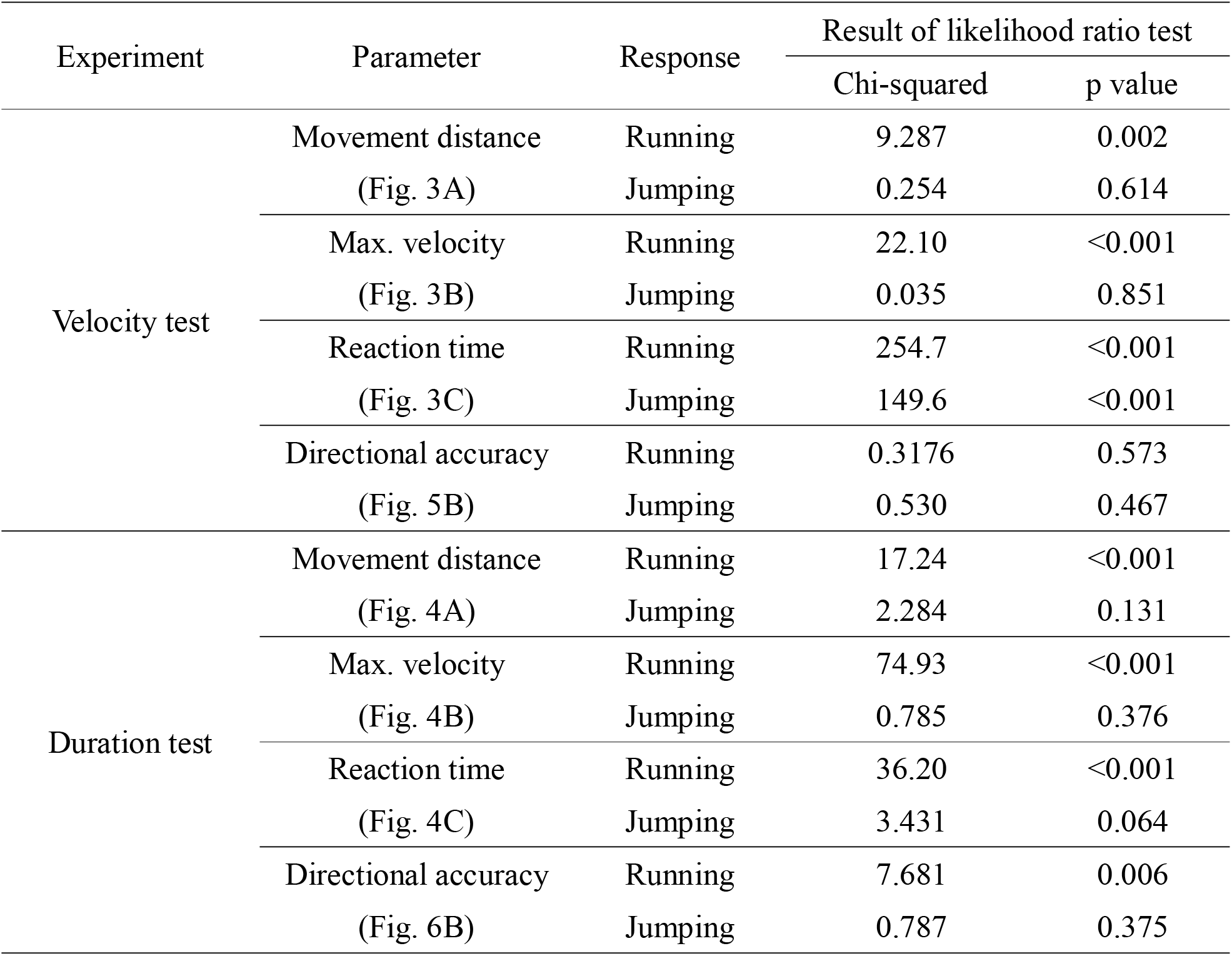
Results of likelihood ratio test for LMMs to test the effects of stimulus velocity and duration on the locomotor parameters.

| Experiment | Parameter | Response | Result of likelihood ratio test |  |
| --- | --- | --- | --- | --- |
|  |  |  | Chi-squared | p value |
| Velocity test | Movement distance<br>(Fig. 3A) | Running | 9.287 | 0.002 |
|  |  | Jumping | 0.254 | 0.614 |
|  | Max. velocity<br>(Fig. 3B) | Running | 22.10 | <0.001 |
|  |  | Jumping | 0.035 | 0.851 |
|  | Reaction time<br>(Fig. 3C) | Running | 254.7 | <0.001 |
|  |  | Jumping | 149.6 | <0.001 |
|  | Directional accuracy<br>(Fig. 5B) | Running | 0.3176 | 0.573 |
|  |  | Jumping | 0.530 | 0.467 |
| Duration test | Movement distance<br>(Fig. 4A) | Running | 17.24 | <0.001 |
|  |  | Jumping | 2.284 | 0.131 |
|  | Max. velocity<br>(Fig. 4B) | Running | 74.93 | <0.001 |
|  |  | Jumping | 0.785 | 0.376 |
|  | Reaction time<br>(Fig. 4C) | Running | 36.20 | <0.001 |
|  |  | Jumping | 3.431 | 0.064 |
|  | Directional accuracy<br>(Fig. 6B) | Running | 7.681 | 0.006 |
|  |  | Jumping | 0.787 | 0.375 |

Relationships between angular parameters and stimulus angle were analyzed by the regression analyses, which were used in our previous study (Sato et al., 2019). The movement direction was considered as non-circular data and linear regression analysis was used as follows:

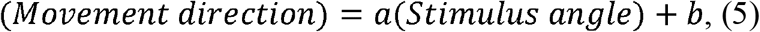

where *a* is the estimated coefficient and *b* is the intercept (Table 4). In contrast, the turn angle was considered as circular data, and circular regression analysis was used as follows:

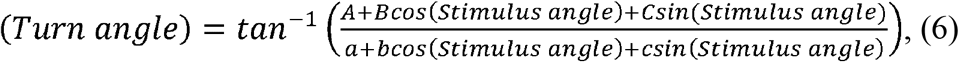

where *A* and *a* are the intercepts, *B* and *b* are the estimated coefficients of cosine, and *c* and *c* are the estimated coefficients of sine in the numerator and denominator of the model, respectively.

**Table 4.**
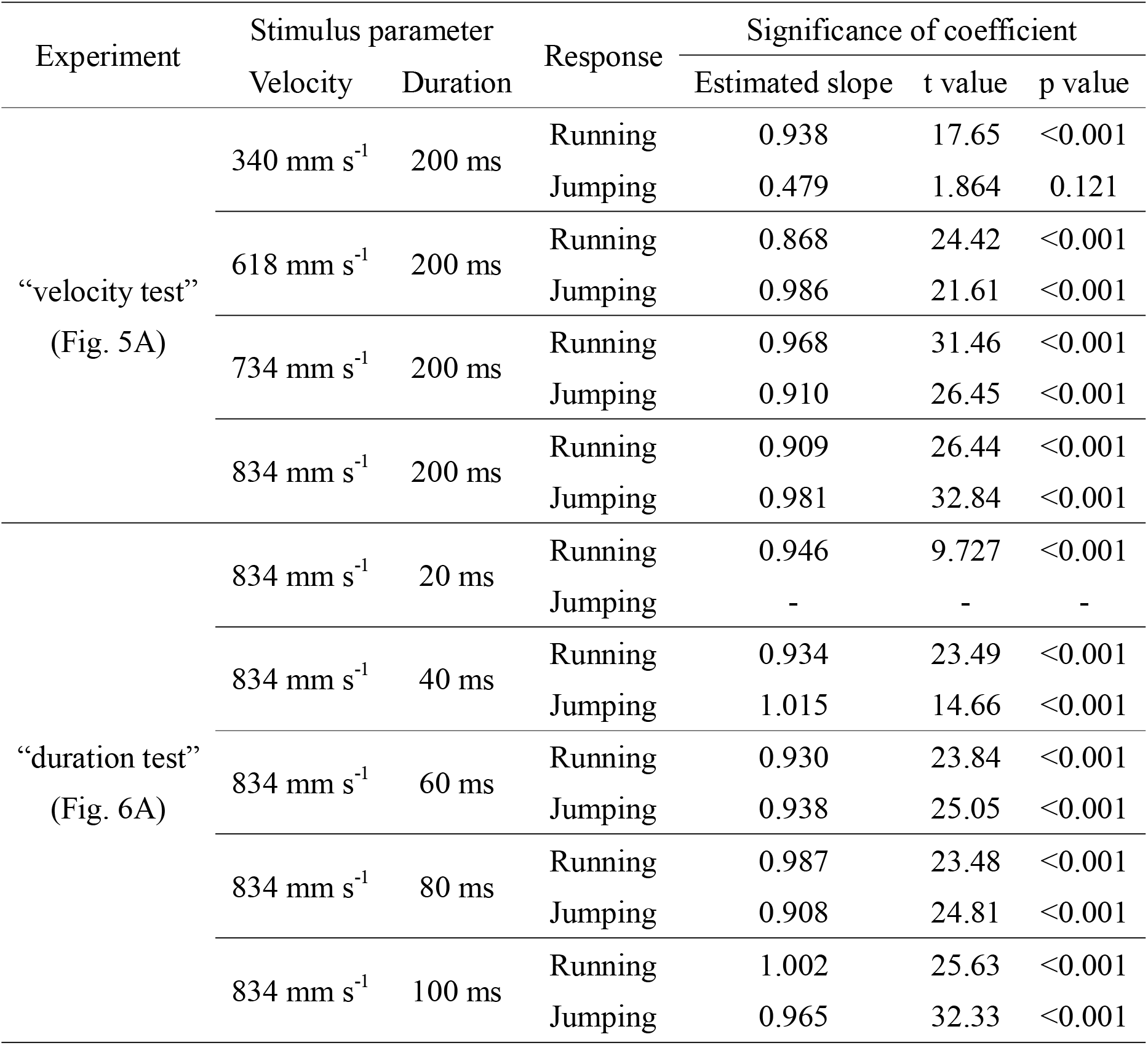
Summary of the result of linear regression analysis for the movement direction and stimulus angle in “velocity test” and “duration test” experiments.

## RESULTS

### Faster and longer stimulus elicited the jumping response more frequently

First, we examined the parameters of the stimulus that affected the action selection of either running or jumping. We tested the effects of angle, velocity, and duration of the stimulus, and found that the crickets chose to jump more frequently in response to the faster and longer stimuli (Fig. 1; Table 1). In the angle test (Fig. 1A,B), the crickets likely chose jumping rather than running in response to the stimulus from behind (approximately 0 degree), but the effect of stimulus angle on the selection ratio of running or jumping was not statistically significant (Table 1). In contrast, the velocity and duration tests demonstrated that the velocity and duration of the stimulus apparently affected both probabilities of the escape response to the stimulus and the selection ratio of running and jumping while escaping (Fig. 1C–F). The response probability and the ratio of jumping choice to running choice significantly increased as the velocity or duration of the stimulus increased (Table 1). These results indicated that the cricket’s choice of escape responses was affected by the two stimulus parameters, velocity and duration of the air current. Although we repeated multiple trials for each individual, the trial orders did not affect the response probabilities or selection ratios in all three experiments (Fig. 1; Table 1).

### Crickets changed running speed depending on stimulus parameters

Based on the data from the angle, velocity, and duration tests, we investigated the effects of stimulus parameters on locomotion performance indicated by metric locomotor parameters. These included movement distance, maximum velocity, and reaction time, in running and jumping. The relationships between these parameters and the stimulus angle indicated that the crickets changed their locomotion performance according to the stimulus angle during running rather than during jumping (Fig. 2). The stimulus angle had a significant impact on all three metric locomotor parameters during running but only on the maximum velocity during jumping (lines in Fig. 2; Table 2). The closer to behind of the cricket the stimulus was applied from, the quicker the crickets began to respond and the farther and faster they ran (left in Fig. 2A–C). In contrast, the closer to front of the cricket the stimulus angle, the faster the crickets jumped, but the movement distance and reaction time of jumping were not significantly affected by the stimulus angle (right in Fig. 2A–C).

**Fig. 2.**
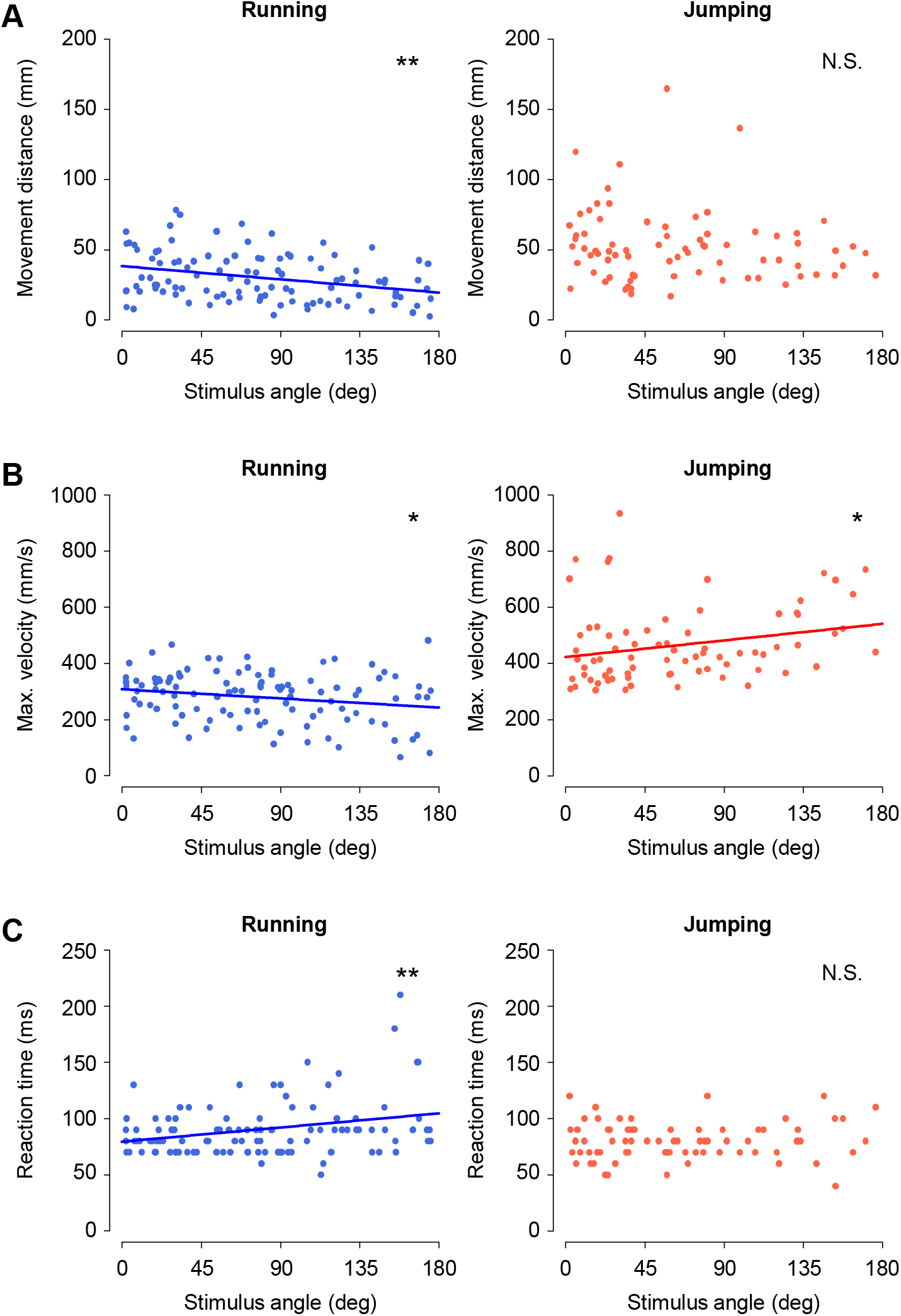
Effects of stimulus angle on metric locomotor parameters. **(A–C)** Relationships between the stimulus angle and the movement distance (A), maximum translational velocity (B), or reaction time (C), in running (left) and jumping (right) in the angle test. The velocity and duration of air current were fixed to 834 mm s^−1^ and 200 ms, respectively. The total number of responses used for the analysis were 106 and 76 for running and jumping, respectively. Lines indicate the regression lines with a significant slope. \**P*<0.05, \*\**P*<0.01, N.S. not significant, linear regression analysis. *N*=10 animals.

In the velocity and duration tests, we examined the effects of velocity or duration of the stimulus not only on the metric locomotor parameters (Figs. 3 and 4) but also on the movement direction (Figs. 5 and 6) of running and jumping. The stimulus velocity and duration influenced the metric locomotor parameters of running rather than jumping. The velocity test indicated that the crickets increased the movement distance and speed during running, but not during jumping, as the stimulus velocity increased (Fig. 3A,B; Table 3). The reaction time became shorter as the stimulus velocity increased in both running and jumping, meaning that the faster the air current, the quicker the crickets initiated the escape regardless of the response type (Fig. 3C; Table 3). In the duration test, the longer the stimulus, the farther and faster the cricket ran, but they did not change their jumping locomotion with the stimulus duration (Fig. 4). The effects of the stimulus duration were significant on all of the metric locomotor parameters during running, whereas none of the parameters in jumping (Table 3). Interestingly, the longer the stimulus, the earlier the cricket started the escape running but did not change the start of jumping. This was probably because the shortest stimulus (20 ms) elicited running responses with a long latency of over 100 ms after the stimulus was terminated (left in Fig. 4C). If the stimulus is too short, it may take longer for the crickets to notice it, or such a stimulus would be judged as not dangerous.

**Fig. 3.**
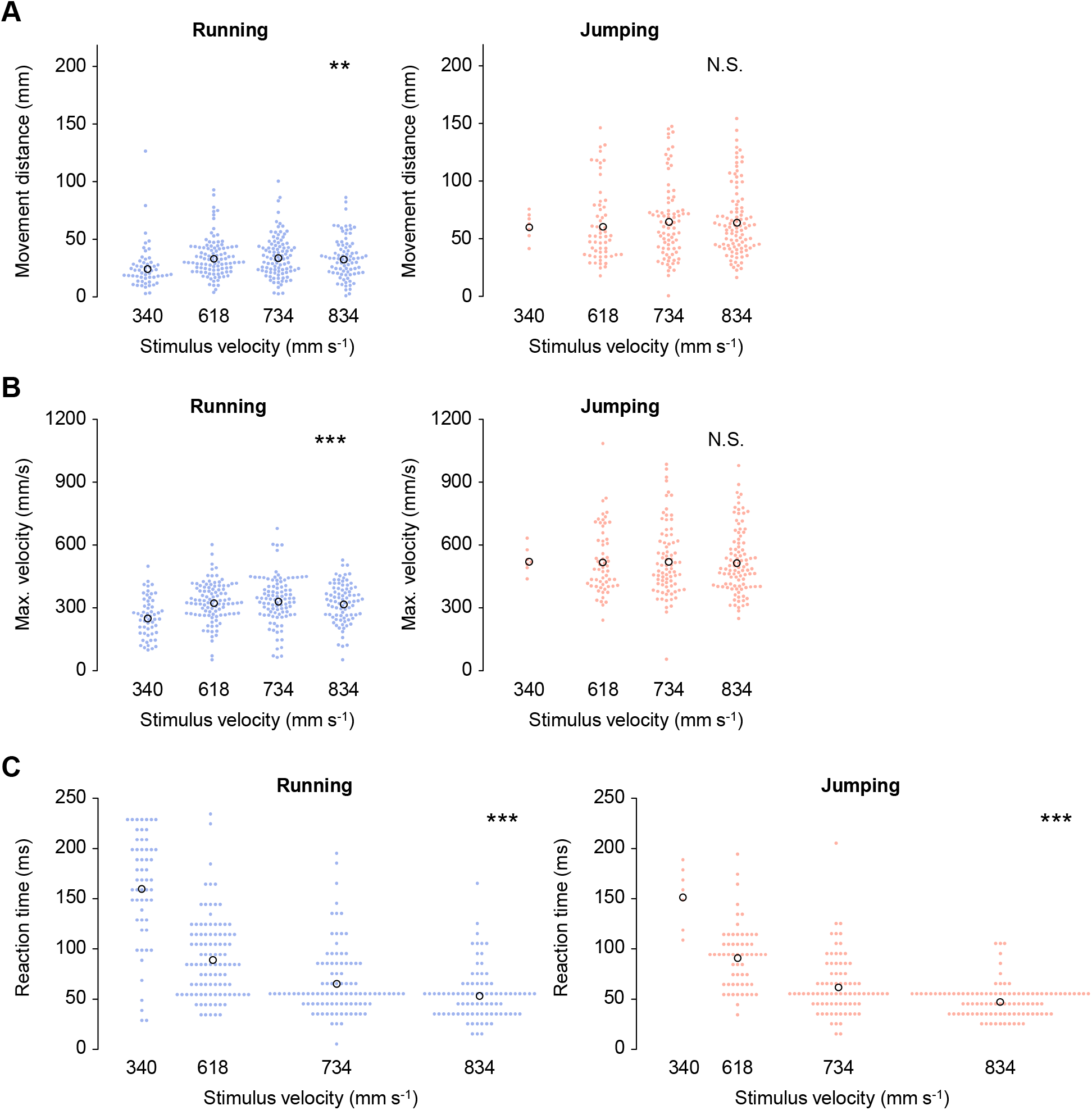
Effects of stimulus velocity on metric locomotor parameters. **(A–C)** The movement distance (A), maximum translational velocity (B), and reaction time (C), in running (left) and jumping (right). The total number of responses used for the analysis were 56, 101, 98 and 84 for running, and 7, 51, 81 and 100 for jumping, for 340, 618, 734 and 834 mm s^−1^ for stimulus velocities, respectively. Colored filled circles represent the data for each trial and black open circles represent the mean of the data in all trials for each velocity of stimuli. \*\**P*<0.01, \*\*\**P*<0.001, N.S. not significant, likelihood ratio test for LMMs. *N*=10 animals.

**Fig. 4.**
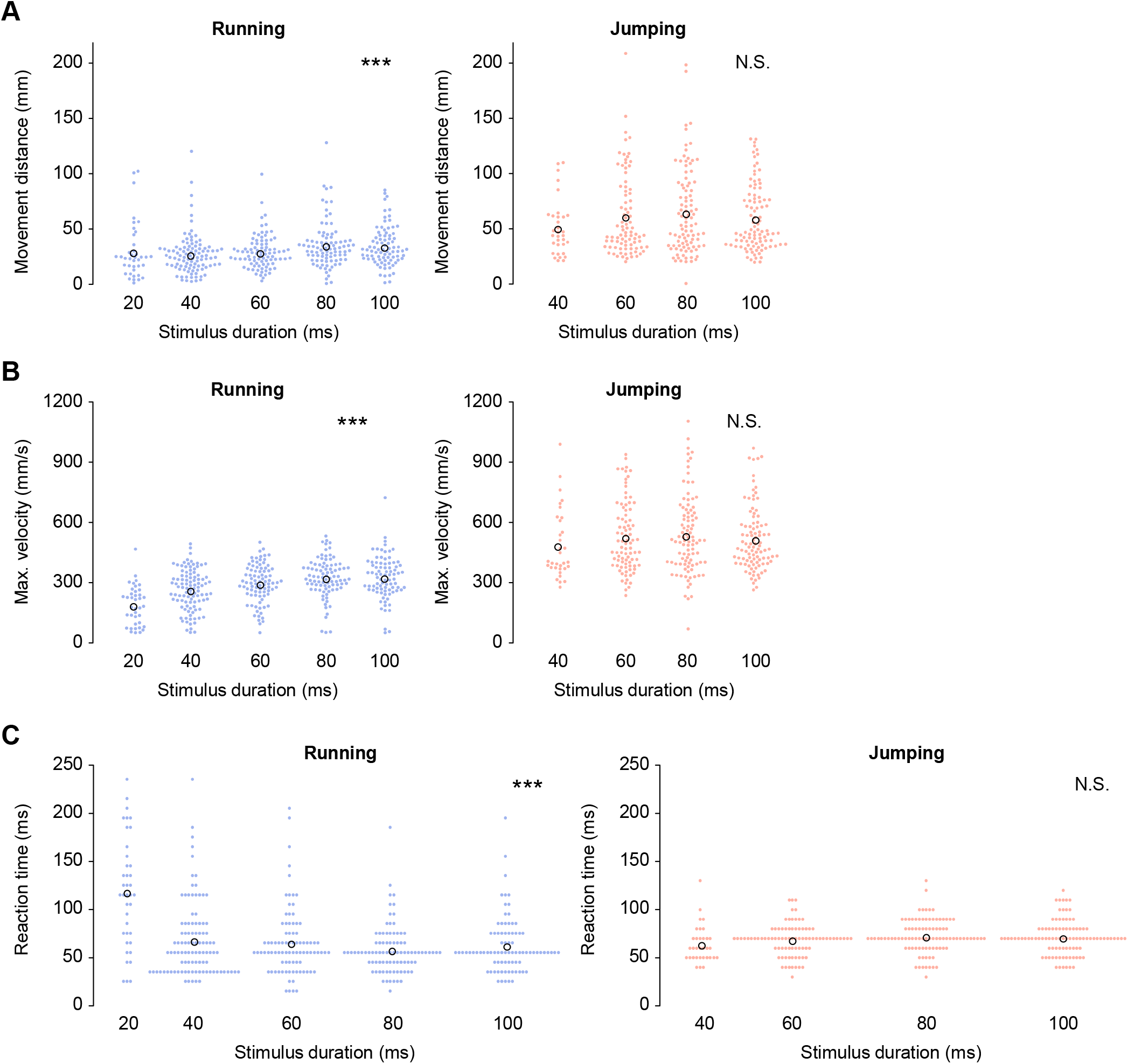
Effects of stimulus duration on metric locomotor parameters. **(A–C)** The movement distance (A), maximum translational velocity (B), and reaction time (C), in running (left) and jumping (right). The total number of responses used for the analysis were 39, 101, 88, 90 and 85 of running, and 0, 33, 87, 97 and 98 of jumping, for 20, 40, 60, 80 and 100 ms of stimulus durations, respectively. Colored filled circles represent the data for each trial and black open circles represent the mean of the data in all trials for each duration of stimuli. No jumping was elicited by the stimulus of 20-ms duration. \*\*\**P*<0.001, N.S. not significant, likelihood ratio test for LMMs. *N*=10 animals.

**Fig. 5.**
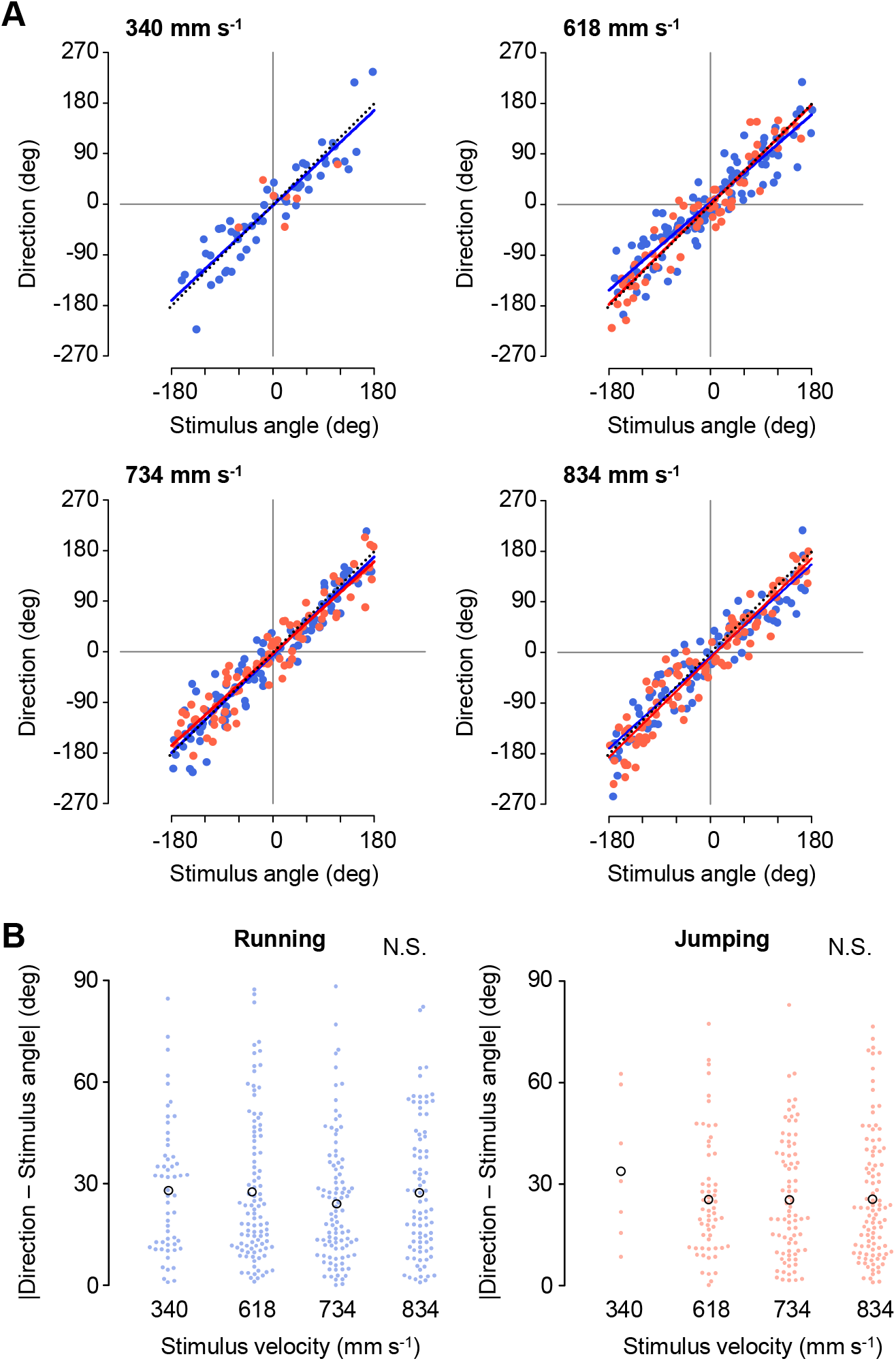
Effects of stimulus velocity on directional control. **(A)** Relationships between the movement direction and stimulus angle in running (blue) and jumping (red), for the stimuli of different velocities, 340, 618, 734, and 834 mm s^−1^. Colored lines represent linear regression lines with a significant slope for the data of running (blue) and jumping (red), respectively. Black dotted lines indicate the line of *y* = *x*. **(B)** Absolute values of the angular difference between the movement direction and stimulus angle in running (left) and jumping (right). Colored filled circles represent the data for each trial and black open circles represent the mean of the data in all trials, for each velocity of stimuli. N.S. not significant, likelihood ratio test for LMMs. *N*=10 animals.

**Fig. 6.**
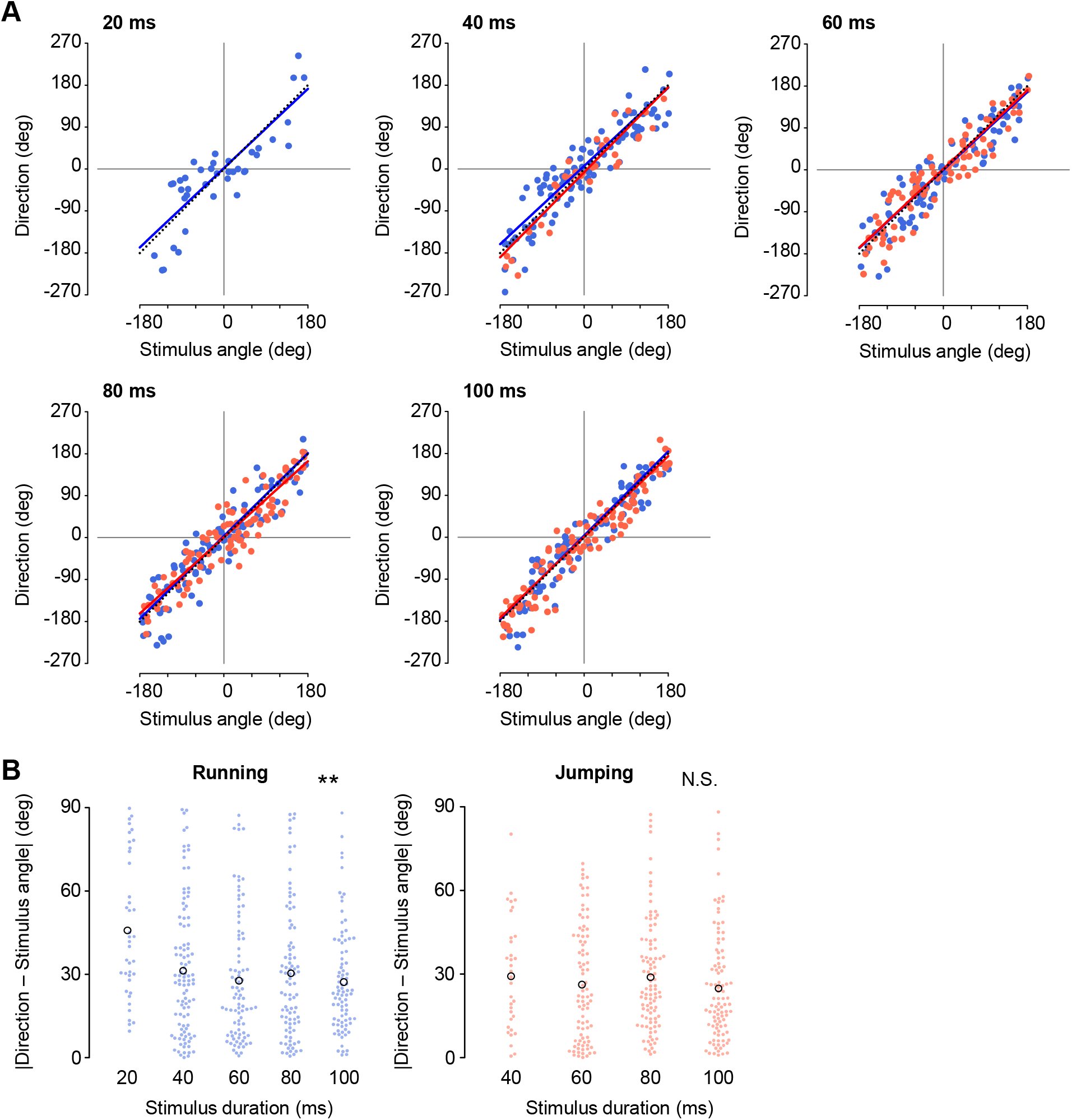
Effects of stimulus duration on directional control. **(A)** Relationships between the movement direction and stimulus angle in running (blue) and jumping (red), for the stimuli of different durations, 20, 40, 60, 80, and 100 ms. No jumping was elicited by the 20 ms duration stimulus. Colored lines represent linear regression lines with a significant slope for the data of running (blue) and jumping (red), respectively. Black dotted lines indicate the line of *y* = *x*. **(B)** Absolute values of the angular difference between movement direction and stimulus angle in running (left) and jumping (right). Colored filled circles represent the data for each trial and black open circles represent the mean of the data in all trials, for each duration of stimuli. \*\**P*<0.01, N.S. not significant, likelihood ratio test for LMMs. *N*=10 animals.

### Movement direction was controlled regardless of stimulus velocity and duration

Our previous studies have shown that crickets precisely control their movement direction against the stimulus angle during both running and jumping (Sato et al., 2019). We then examined whether the velocity and duration of the stimulus affected the directional control of the cricket escape responses (Figs. 5 and 6). The crickets could control their movement direction precisely against the stimulus angle, independent of the velocity and duration of the stimulus. Even in response to stimuli of different velocities or durations, plots of the movement direction against stimulus angle were distributed close to the *y* = x line during running and jumping (Figs. 5A and 6A; Table 4). Here, how precisely the crickets moved in the opposite direction to the stimulus was assessed by the absolute value of the angular difference between the moving direction and stimulus angle. Independent of the stimulus velocity, the absolute angular differences between the movement direction and stimulus angle were approximately 30 degree in both running and jumping (Fig. 5B; Table 3), indicating no effect of the air current velocity on the directional control. Also, in the duration test, the duration of the stimulus had little impact on the directional control of the escape responses (Fig. 6). The absolute angular difference between the movement direction and stimulus angle was mostly constant for the stimuli of 40–100 ms duration both in running and jumping (Fig. 6B). The significant effect of the stimulus duration was observed in running but not in jumping (Table 3), which may be because the running directions in response to the shortest (20 ms) stimulus were less correlated with the stimulus angle. In contrast, no jumping was elicited by the shortest stimuli, resulting in no significant difference in the absolute difference value of the stimulus duration in jumping. We also checked the impact of the velocity and duration of the stimulus on the relationship between the turn angle and stimulus angle (Figs. S2 and S3) but observed no apparent difference in the distributions of the plot among the responses either in the velocity test or in the duration test. Thus, the angular parameters of escape responses, including the movement direction and turn angle, were not significantly affected by the velocity and duration of the stimulus, except for extremely short stimuli that were close to the threshold to induce jump.

To conclude regarding the impacts of stimulus parameters on locomotor parameters, crickets altered their locomotion performance, such as the moving distance, velocity, and reaction time, according to the angle, velocity, and duration of the stimulus during running but hardly during jumping. In contrast, the escape direction was controlled precisely, independent of the velocity and duration of the stimulus in both running and jumping escape responses. These results suggest that movement performance in speed and distance is modulated during the running response but not while jumping, which depends on the stimulus characteristics. Directionality of the escape movements was dictated by the stimulus angle only, but was unaffected by the stimulus intensity and duration.

## DISCUSSION

### The neural system to choose escape response is based on multiple stimulus parameters

The prey animal must perceive how the predator approaches through multiple aspects of the stimulus in order to perform the most successful escape response. Previously, the choice of escape response has been reported to depend on stimulus velocity (Bhattacharyya et al., 2017; von Reyn et al., 2014, 2017). Our results revealed that both velocity and duration of the stimulus affect the action selection of either running or jumping in the wind-elicited escape behavior of crickets. In conclusion, crickets likely made a decision regarding their escape responses based on multiple aspects of the air-current stimuli that indicate the predator’s approaches. This finding strongly suggests that there is an integration process of multiple channels of distinct stimulus information for action selection in the crickets’ central nervous system.

While complex neural mechanisms across multiple brain regions are involved in mammalian escape behaviors (Evans et al., 2018; Gross and Canteras, 2012; Shang et al., 2015; Wang et al., 2015), relatively simple neural circuits mediate the escape behavior in fish and invertebrates (Card, 2012; Eaton et al., 2001). In such simple neural systems that mediate escape behavior, a few neurons have been identified that trigger the escape responses to a specific sensory stimulus which indicate the predator’s approach. In flies, a key set of neurons called giant fibers directly induce quick taking-off as an escape response (Allen et al., 2006). To choose from two types of escaping take-off depending on the stimulus velocity, the presence of neural circuits in which specific projection neurons provide information of stimulus velocity to the giant fiber have been proposed in the fly (von Reyn et al., 2014, 2017). In context, our results show that crickets choose escape responses based on multiple stimulus parameters for the first time, which would promote an understanding of the mechanism of choosing appropriate behavior according to varied situations in the natural field.

The neural basis of the cercal sensory system to process airflow information, which in turn mediates wind-elicited escape behavior, has been well studied (Jacobs et al., 2008; Baba and Ogawa, 2017). Air currents are detected as air-particle displacement by filiform hairs on the cerci (Miller et al., 2011; Shimozawa and Kanou, 1984), and several wind-sensitive interneurons, including giant interneurons (GIs), which have been identified as ascending projection neurons, receive excitatory synaptic inputs from the sensory afferents of the mechanoreceptor neurons of the hair sensilla. The GIs encode various characteristics of the surrounding air currents such as velocity, direction, and frequency in their firing activity (Aldworth et al., 2011; Miller et al., 1991), and project their long axons through the ventral nerve cord to higher centers, including the thoracic ganglia and the brain (Hirota et al., 1993). Because the GIs differs from each other in their sensitivities to the direction and intensity of the air currents (Jacobs et al., 2008; Miller et al., 1991; Ogawa et al., 2008), the GIs likely provide distinct sensory information to their postsynaptic partners. Thus, further investigation of the postsynaptic circuit of GIs would clarify the neural mechanism underlying action selection depending on the multiple stimulus parameters.

### Stimulus characteristics that were likely to elicit jumping

The faster and longer the air currents, the more frequently the jumping response was elicited. This is due to the velocity and duration of the air-current stimulus being considered to be correlated with the size of the predator and their approaching speed (Casas and Dangles, 2010), crickets would choose to jump in situations where a larger predator attacked them more quickly. It should be noted, however, that the jumping probability was saturated at approximately 50% even for the fastest and longest stimulus, which was the most inducible for the jump (Fig. 1). One possibility is that additional sensory inputs may be required to further elevate the likelihood of jumping selection. Animals use a variety of sensory modalities to detect predators. For example, auditory stimuli combined with air currents alter the contents of escape responses in crickets (Fukutomi and Ogawa, 2017; Fukutomi et al., 2015). Even if they do not directly elicit escape responses, other sensory inputs that inform external contexts or internal conditions of the animals affect the escape responses (Domenici et al, 2008; Matsuura et al., 2002). Therefore, crickets may jump more frequently than 50% in different contexts or situations.

In contrast, the stimulus angle did not significantly affect the selection of running and jumping. This result is consistent with a previous report where crickets have the same accuracy of directional controllability during running and jumping (Sato et al., 2019). In this study, a strong correlation, which could be approximated by *y* = *x*, between the movement direction and stimulus angle, was also observed in the velocity test (Fig. 5) and the duration test (Fig. 6). These results indicated that the escape direction could be controlled against the stimulus angle in either type of escape response, independent of the stimulus velocity and duration. It is supposed that the sensory information of the stimulus angle would be conveyed by neural channels different from those for stimulus velocity and duration.

### Stimulus dependency of the locomotion performance

The crickets changed their moving speed and travel distance according to the stimulus parameters during running, but not during jumping. This locomotion controllability is the behavioral advantage of running compared to jumping. In contrast, the invariance of the jumping speed and distance to the stimulus parameters illustrated that jumping was a more stereotyped response, which might be more predictable for predators. These results are consistent with our previous reports that running is a more flexible response during which crickets respond to the additional stimulus, compared to jumping (Sato et al., 2019).

Interestingly, the reaction time decreased as the stimulus velocity increased during both running and jumping (Fig. 3C). Previous studies have demonstrated that the escape responses to visual looming stimuli start at a specific angular size of the stimulus, which indicates the distance to the stimulus. This suggests that the distance to the predators, rather than the speed of the predator’s approach, is a more crucial sensory cue for the prey to decide an escape start (Bhattacharyya et al., 2017; Dunn et al., 2016; Fotowat and Gabbiani, 2007). In contrast, our results revealed a strong relationship between stimulus velocity and escape latency. This means that the crickets may decide to start escaping based on the stimulus velocity, which indicates the approaching speed of a predator. The faster a predator approaches, the less time it takes to reach its prey; thus, the reaction time is important for the prey to increase the success rate of the escape behavior (von Reyn et al., 2014; Walker et al., 2005). In particular, when animals of prey cannot visually detect the predator’s approach, as in the dark, they will make decisions regarding the escape start based on the stimulus velocity for effective survival.

Unlike the metric locomotor parameters, the movement direction was accurately controlled almost independent of velocity or duration of the stimulus. This result was consistent with a previous study reporting that the direction of the escape running on a treadmill is similarly controlled to stimuli of different durations (Oe and Ogawa, 2013). The accuracy of directional control decreased only in the running response to the shortest stimulus (left in Fig. 6B). There is a possibility that a 20 ms stimulus is too short for the crickets to perceive their direction. This is also supported by the result where the reaction time for running response to the shortest stimulus was longer than that for stimuli with longer durations (left in Fig. 4C). Air currents longer than a certain duration will likely be necessary for the cricket to perceive stimulus orientation for directional control of the escape response.

## Acknowledgements

We thank Dr Masayo Soma for helpful advice regarding statistical analysis.

## Author contributions

Conceptualization: N.S., H.S., H.O.; Methodology: N.S., H.S., H.O.; Validation: N.S.; Formal analysis: N.S.; Investigation: N.S.; Data curation: N.S.; Writing – original draft: N.S.; Writing – review & editing: H.S. and H.O.; Visualization: N.S.; Supervision: H.O.; Project administration: N.S., H.S., H.O.; Funding acquisition: N.S. and H.O.

